# Charting the human amygdala development across childhood and adolescence: Manual and automatic segmentation

**DOI:** 10.1101/2021.02.11.430883

**Authors:** Quan Zhou, Siman Liu, Chao Jiang, Ye He, Xi-Nian Zuo

## Abstract

The developmental pattern of the amygdala throughout childhood and adolescence has been inconsistently reported in previous neuroimaging studies. Given the relatively small size of the amygdala on full brain MRI scans, discrepancies may be partly due to methodological differences in amygdalar segmentation. To investigate the impact of volume extraction methods on amygdala volume, we compared *FreeSurfer, FSL* and *volBrain* segmentation measurements with those obtained by manual tracing. The manual tracing method, which we used as the ‘gold standard’, exhibited almost perfect intra- and inter-rater reliability. We observed systematic differences in amygdala volumes between automatic (*FreeSurfer* and *volBrain*) and manual methods. Specifically, compared with the manual tracing, *FreeSurfer* estimated larger amygdalae, and *volBrain* produced smaller amygdalae while *FSL* demonstrated a mixed pattern. The tracing bias was not uniform, but higher for smaller amygdalae. We further modeled amygdalar growth curves using accelerated longitudinal cohort data from the Chinese Color Nest Project. Trajectory modeling and statistical assessments of the manually traced amygdalae revealed linearly increasing and parallel developmental patterns for both girls and boys, although the amygdalae of boys were larger than those of girls. Compared to these trajectories, the shapes of developmental curves were similar when using the *volBrain* derived volumes. *FreeSurfer* derived trajectories had more nonlinearities and appeared flatter. *FSL* derived trajectories demonstrated an inverted U shape and were significantly different from those derived from manual tracing method. The use of amygdala volumes adjusted for total gray-matter volumes, but not intracranial volumes, resolved the shape discrepancies and led to reproducible growth curves curves between manual tracing and the automatic methods (except *FSL*). Our findings revealed steady growth of the human amygdala, mirroring its functional development across the school age. Methodological improvements are warranted for current automatic tools to achieve more accurate tracing of the amygdala at school age, calling for next generation tools.

## 1. Introduction

Childhood and adolescence are key periods for socioemotional development, which correlate strongly with the development of risk factors for diverse neuropsychiatric disorders (Paus et al., 2008). Together with enhanced efforts to prevent such disorders, many large-scale studies have been undertaken to explore behavioral and biological development of children and adolescents (Ortiz and Raine, 2004; Silk et al., 2007; Connor, 2004). Rapid progress in in-vivo brain imaging technologies has accelerated the use of structural magnetic resonance imaging (MRI) to quantify volumes of different brain structures. These morphological features have been demonstrated by MRI to be sensitive for developmental brain changes (Tamnes et al., 2013; Albaugh et al., 2017; Wierenga et al., 2018). The accurate developmental trajectories of brain structures using MRI is thus an important requirement for understanding the neurodevelopmental mechanisms of these disorders occurring during childhood and adolescence.

The amygdala is an almond-shaped brain structure, part of the limbic system and is highly connected with other brain regions (Schumann and Amaral, 2005). It plays important roles in emotional and cognitive processes, especially fear and threat processing (LeDoux, 1998; Cardinal et al., 2002; Pessoa, 2010) and exhibits network-level connectivity changes across the human lifespan (He et al., 2016). Abnormal amygdalar structure in children and adolescents has been related to a plethora of neurodevelopmental abnormalities (Scherf et al., 2013; Schumann et al., 2011), including autism (Mosconi et al., 2009; Schumann et al., 2004), anxiety disorder (De Bellis et al., 2000; Redlich et al., 2015) and schizophrenia (Ganzola et al., 2014). Meanwhile, many studies have explored age-related changes of the amygdala in pediatric and adolescent samples (Uematsu et al., 2012; Gilmore et al., 2012; Wierenga et al., 2014; Barnea-Goraly et al., 2014; Herting et al., 2018), indicating the promise of using normal growth patterns for monitoring abnormal development. Growth charts are expected to aid risk evaluation, early diagnosis and educational monitoring by delineating typical development standards. In several recent studies, researchers have tracked the age-related increase of amygdala volume from childhood through adolescence (Herting et al., 2018; Goddings et al., 2014; Albaugh et al., 2017). However, a study including 271 individuals aged 8-29 years reported no significant changes in amygdala volume (Wierenga et al., 2018). This was similar to the observation from a sample of 85 individuals scanned twice across 8-22 years (Tamnes et al., 2013). Thus, there are mixed findings in the literature related to age-related differences or changes in amygdala volume. The anatomical complexity can limit the accurate measurement of amygdalar volume, leading to a large variation in findings obtained using different amygdala segmentation methods (Lyden et al., 2016), which may explain this inconsistency and less reproducibility (Mills and Tamnes, 2014; Lyden et al., 2016).

Manual tracing is commonly considered the ‘gold standard’ for amygdala segmentation (Morey et al., 2009). It enables flexible quantification guided by prior anatomical knowledge, without the need to make any of the assumptions built into algorithms. Experienced human tracers can correctly label ambiguous borders by adjusting for variation caused by complex or atypical anatomy and image artifacts. To increase reliability and reduce potential biases associated with manual tracing, multiple protocols have been generated and described in the literature (Schumann et al., 2004; Pruessner et al., 2000; Watson et al., 1992). These protocols significantly increase intra- and inter-rater agreement (Pruessner et al., 2000). However, manual tracing is time-consuming and requires the operator to have sufficient anatomical expertise. For large MRI datasets, the labor cost of manual tracing is prohibitive (Akudjedu et al., 2018; Schmidt et al., 2018). There is also subtly drift in tracing criteria of manual raters during the course of a long study. Accordingly, it is critical to develop automatic techniques that can accurately segment amygdala structures from large and growing datasets while providing consistent results and minimizing the human effort necessary for manual tracing.

Several tools have been developed to achieve automatic segmentation in a time-efficient manner including *FSL*-FIRST, *FreeSurfer* and *volBrain*, which are both freely available, ease to use, nearly fully automated, and very accurate (Fis-chl et al., 2002; Manjón and Coupé, 2016; Morey et al., 2009; Akudjedu et al., 2018; Schmidt et al., 2018; Næss-Schmidt et al., 2016). FIRST was provided as part of the *FSL* software library to estimate boundaries of brain structures based on the signal intensity of the T1 image as well as the expected shape of structures using a probabilistic framework (Patenaude et al., 2011). *FreeSurfer* automatically assigns a label to each voxel from anatomical images based on probabilistic estimations relying on Markov random fields (Fischl et al., 2002). It may be difficult for this model-based method applied in *FreeSurfer* to model the regions of interest with sufficient accuracy in highly variable MRI data, such as inter-individual differences or pathological changes in neuroanatomy. To address this, multi-atlas label fusion approach such as *volBrain* has also been implemented. Multi-atlas label fusion segmentation techniques could combine multiple atlas information, thereby minimizing mislabeling from inaccurate affine or non-linear registration (Manjón and Coupé, 2016). Although automated segmentation has been shown to be comparable to manual tracing for adult populations (Fischl et al., 2002; Manjón and Coupé, 2016; Morey et al., 2009; Grimm et al., 2015), its performance for child and adolescent samples, in which head size and shape as well as the pace of structural growth differ, has not been validated adequately (Herten et al., 2019). In addition, the effects of any differences in the accuracy of automatic and manual amygdala segmentation on the subsequent examination of amygdala development in school-age children and adolescents remain incompletely understood.

To fully characterize similarities and discrepancies among techniques, we compared amygdala volumes obtained manually to those extracted by *FreeSurfer, volBrain* and FIRST in *FSL* using 427 longitudinal structural MRI scans from 198 healthy children and adolescents (baseline age: 6-17 years). To answer the aforementioned question, we examined how different tracing methods lead to trajectory differences in amygdala development across school age. Based upon previous reports (Morey et al., 2009; Schoemaker et al., 2016), we expected to observe systematic differences in amygdala segmentation performance among the three tracing methods. We hypothesized that such differences would affect the modeling of human amygdala growth.

## 2. Materials and Methods

### 2.1. Participants

The sample described in this study was part of an accelerated longitudinal database, namely the Chinese Color Nest Project (CCNP: http://deepneuro.bnu.edu.cn/?p=163) for developmental brain-mind association studies across different stages of the postnatal lifespan (Zuo et al., 2017; Liu et al., 2020). Such acceleration was implemented by mixing cross-sectional and longitudinal design to achieve long-time follow-up studies, such as lifespan development cohorts (Nooner et al., 2012; Thompson et al., 2011). It was part of the developmental component of CCNP (devCCNP), and collected at Southwest University (devCCNP-SWU), Chongqing, China. The devCCNP was designed to delineate normative trajectories of brain development in the Chinese population across the school-aged years. The participants had no neurological or mental health problem and did not use psychotropic medication; their estimated intelligence quotients were ⩾ 80. The devCCNP-SWU samples included data from 201 typically developing controls (TDCs) aged 6-17 years who were invited to participate in three consecutive waves of data collection at intervals of approximately 1.25 years (Dong et al., 2020, 2021). T1-weighted MRI examinations were performed at these time points, and the images were visually inspected to exclude those with substantial head-motion artifacts and those with structural abnormalities. After this initial quality control, the final sample included 427 scans from 198 participants (105 females; 93 males; **Table 1**). Scans from three time points, two time points, and one time point were available for 79, 71, and 48 participants, respectively. The mean number of scans per participant was 2.16 (standard deviation = 0.79). The current study was approved by review committees of the participating institutions (Institute of Psychology, Chinese Academy of Sciences, and Southwest University).

### 2.2. MRI acquisition

All participants underwent MRI examinations performed with a Siemens Trio™ 3.0 Tesla MRI scanner. A high-resolution magnetization-prepared rapid gradient-echo (MP-RAGE) T1 sequence (matrix = 256 × 256, FOV = 256 × 256 mm^2^, slices thickness = 1mm, repetition time (TR) = 2600 ms, echo time (TE) = 3.02 ms, inversion time (TI) = 900 ms, flip angle = 15°, number of slices = 176) was obtained for each individual.

### 2.3. Volumetric MRI preprocessing and segmentation

All the images were anonymized by removing all the personal information from the raw MRI data. We removed the facial information by using the **facemasking** tool (Milchenko and Marcus, 2013). The anonymized images were then uploaded to the online image processing system *volBrain* (http://volbrain.upv.es) for brain extraction (Manjón and Coupé, 2016). All the extracted individual brains were also denoised by spatially adaptive non-local means and corrected for intensity normalization in *volBrain*. These preprocessed brain volumes were all in the native space and ready for subsequent manual and automatic tracing procedures. Of note, we confirmed that the impacts of the face masking on the brain extraction and preprocessing are trifling by checking the individual images.

#### 2.3.1. Manual tracing and reliability assessment

Anatomically trained raters QZ (the first author Quan Zhou) and ZQZ performed manual amygdala segmentation in the native space using the ITK-SNAP software (ver. 3.8.0) (Yushkevich et al., 2006). The anatomical boundaries of amygdala structures were defined and segmented according to the protocol described by Pruessner et al. (2000). This protocol has been demonstrated to achieve almost perfect intra- and inter-rater reliability. The reliability was quantified with intraclass correlation coefficient (ICC), which was interpreted as indicating slight [0, 0.20), fair [0.20, 0.40), moderate [0.40, 0.60), substantial [0.60, 0.80), or almost perfect [0.80, 1] reliability (Landis and Koch, 1977). To assess reliability for the protocol implementation in this study, QZ and ZQZ independently traced the amygdala volumes of 30 scans twice at a two-week interval. They were chosen from 30 subjects at baseline examination balanced for age and sex. The ICCs with a 95% confidence interval (CI) are derived by the following hierarchical linear mixed model on the repeated tracing volumes

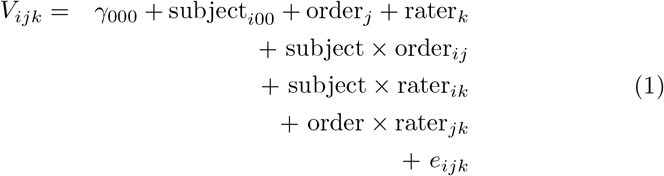

where *V*_*ijk*_ represents the amygdalar volume measurement for the *i*−th (*i* = 1, 2, …, 30) participant in the *j*−th (*j* = 1, 2) manual tracing by the *k*−th rater (*k* = 1, 2); *γ*_000_ is the intercept for a fixed effect of the group average; the following three terms represent random effects for the *i*−th participant, the *j*−th tracing order, the *k*−th rater, respectively; and other three terms denote random interaction effects between the *j*−th tracing and the *i*−th participant, between the *k*−th rater and the *i*−th participant, between the *j*−th tracing and the the *k* −th rater; and *r*_*ijk*_ is an error term.

The above-mentioned model assumes that the seven included variables are independent and distributed normally with zero means. The total variances can be decomposed into the variance component:

- among participants 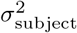
- between repeated tracings by the same rater 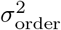
- between raters for the same tracing order 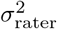
- among participants due to the differences in tracing order 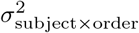
- among participants due to the differences in rater 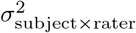
- between two raters due to the differences in tracing order 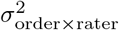
- of the residual 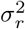.

We define the inter-rater reliability of the human amygdala volumetric measurements by manual tracing as:

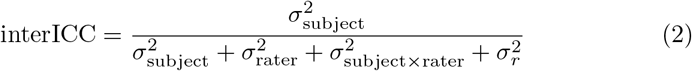

and the intra-rater reliability of the human amygdala volumetric measurements by manual tracing as:

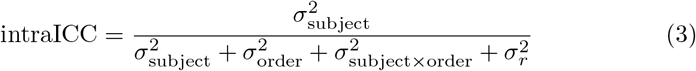

#### 2.3.2. Automatic tracing and visual inspection

Amygdala volumes were estimated using *volBrain* (http://volbrain.upv.es), a fully automated segmentation method that has outperformed other segmentation methods across many brain structures (Manjón and Coupé, 2016). The operational pipeline has been described and evaluated previously (Manjón and Coupé, 2016). Intracranial volume (ICV) and total gray matter volume (GMV) were derived with *volBrain*. Amygdala volumes were also obtained using *FSL*-FIRST (v.6.0.4; http://fsl.fmrib.ox.ac.uk/fsl/fslwiki/FIRST). More detailed information about the processing steps of subcortical segmentation by *FSL*-FIRST can be found in (Patenaude et al., 2011). In the current study, we segmented the T1 images using *FSL*-FIRST with none boundary correction. Amygdala segmentation labels were saved as binary masks. A voxel count was subsequently used to calculate the amygdalar volumes. Both ICV and GMV were also obtained using *FSL. FreeSurfer* segmentation includes the cross-sectional (CS-FS) and the longitudinal streams (LG-FS). Automatic segmentation and labeling of the human amygdala were also performed using the “recon-all” pipeline in CS-FS and LG-FS (ver. 6.0.0; http://surfer.nmr.mgh.harvard.edu). These processing stages have been documented in (Fischl et al., 2002; Reuter et al., 2012). Amygdala volumes provided in **aseg.stats** files were used in the subsequent analysis, and **aseg.mgz** volume files were converted into NIFTI files in native space for visualization. Transformation for segmentation and its inverse transformation to native space for volumetric comparison have been described in (Morey et al., 2009). Both ICV and GMV measurements were also provided by the CS-FS outputs while The GMV measurements were also provided by the LG-FS outputs. Due to LG-FS’s possible bias by matching head across all the time points in children, ICV measurements were not obtained by LG-FS. All segmentation results were visually inspected to ensure adequate registration. Visual inspection of the traced amygdala volumes in a representative subject using manual and automatic methods is illustrated in **Figure 1**.

**Figure 1:**
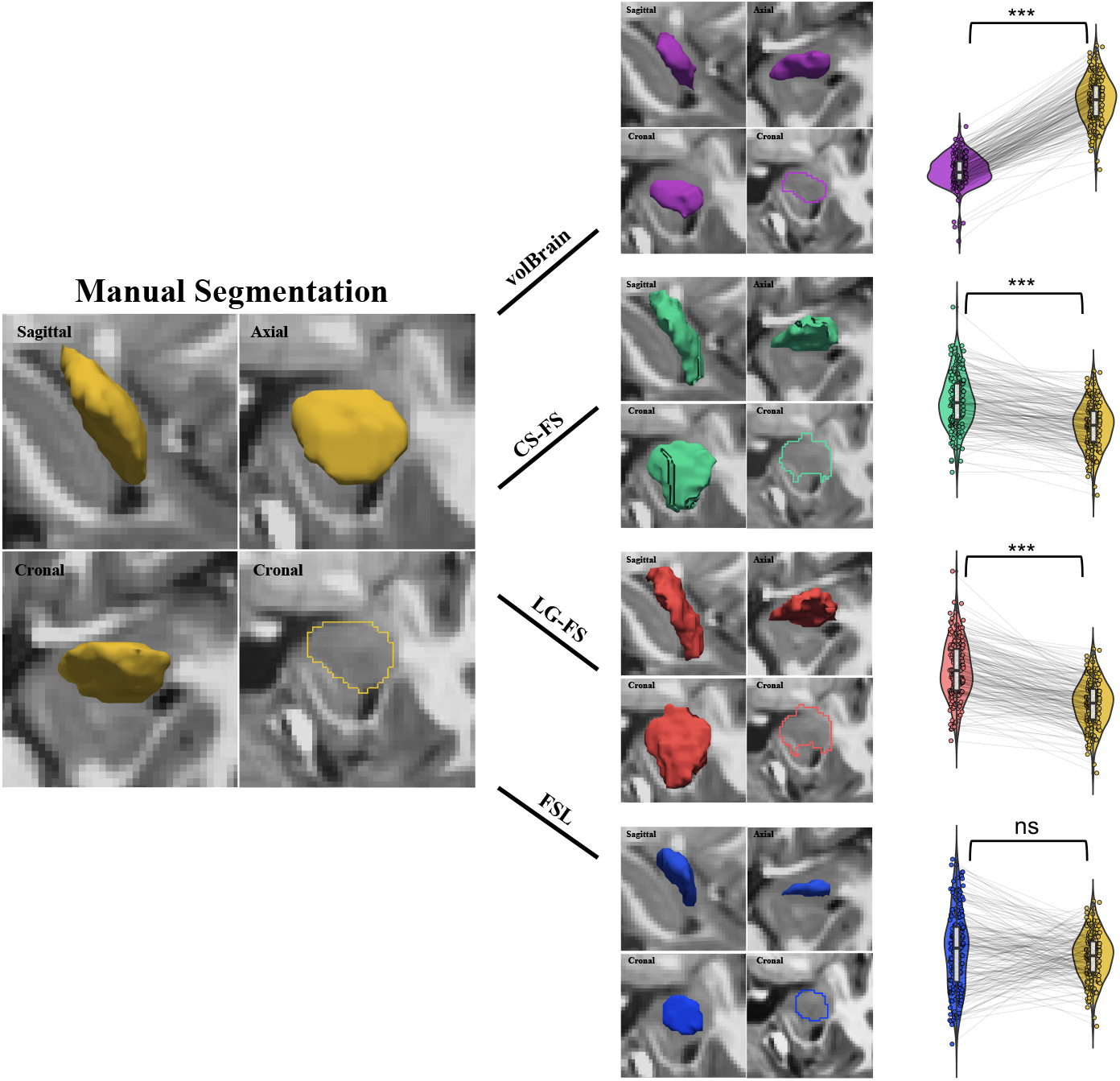
The amygdala structures extracted from five different techniques. A sample participant’s segmented amygdala by manual tracing (Yellow), *volBrain* (Purple), CS-FS (Green), LG-FS (Red) and *FSL* (Blue). Scatter Violin plots present the paired changes of the traced left amygdala volumes between manual tracing and *volBrain*, CS-FS, LG-FS and *FSL* for the first-wave devCCNP samples. ∗*p* < 0.05; ∗ ∗ *p* < 0.01; ∗ ∗ ∗*p* < 0.001

### 2.4. Accuracy assessments on automatic segmentation

QZ manually traced all the 427 amygdala of the devCCNP-SWU samples, which served as the reference volumes (i.e., gold standard) for the subsequent analyses. We validated the accuracy of automatic segmentation separately for each of the three waves of the samples. For each wave, we performed paired t-tests on traced volumes between the automatic and manual methods. We quantified volume difference between the automatic and manual tracing as the equation 4. A greater volume difference indicates increased discrepancy relative to the manually segmented amygdala volumes. To examine systematic changes of the traced volumes, we tested the Pearson’s correlation of traced volumes between the automatic and manual methods across individual subjects. A strong correlation (*R ≥* 0.8) is taken to indicate good consistency on the individual differences in amygdala volumes between the manual and automatic methods. We further calculated the spatially overlapping volumes, the false positive rate and the false negative rate to quantitatively measure the degree of correct or incorrect estimation of the automatic methods. These metrics are defined as:

- percentage of volume difference

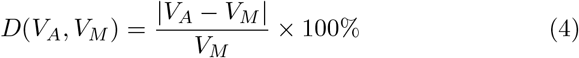
- percentage of spatial overlap

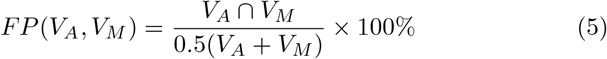
- false-positive rate

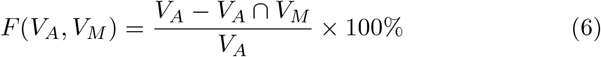
- false-negative rate

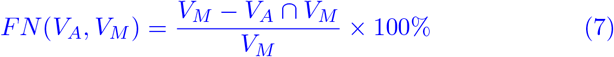

In these equations, *V*_*A*_ is the volume measured automatically and *V*_*M*_ is that measured manually (the reference, i.e., the gold standard). The maximum *P*(*V*_*A*_, *V*_*M*_) value is 100%, reflecting identical tracing between manual and automatic method while smaller values indicated less perfect spatial overlaps (Morey et al., 2009), implying the worse performance of the automatic tracing. The minimum *FP*(*V*_*A*_, *V*_*M*_) value is 0, reflecting identical tracing between manual and automatic method while larger values indicate higher error rates of automatic segmentation, i.e., the inclusion of larger proportions of non-amygdalar structure(s). The minimum *FN*(*V*_*A*_, *V*_*M*_) value is 0, reflecting identical tracing between manual and automatic segmentation while higher values indicate more error rates of automated protocol, i.e., the exclusion of larger proportions of amygdalar structure(s).

To further investigate how the accuracy of the automatic tracing methods varies with amygdala sizes, we employed a generalized additive mixed model (GAMM) to model the size effect of amygdala on the automatic tracing accuracy. In addition, young participants are more likely to move during scan acquisition, leading to worse scan quality and motion artifacts, which may affect the precise differentiation of structures by automated techniques. To exclude the effects of image quality on segmentation accuracy, we included motion and scan quality as covariates for the regression models. Specifically, the motion metric is the coefficient of joint variation (CJV) as an objective function for the optimization of intensity non-uniformity correction algorithms (Ganzetti et al., 2016) while the image quality is quantified by signal-to-noise ratio (SNR). Lower CJV and higher SNR values indicate better image quality (Welvaert and Rosseel, 2013). Specifically, we plotted the spatial overlap (the overlap percentage *P*) between automatic and manual segmentation as a function of the reference (i.e., the manually traced) volumes, while controlling CJV and SNR. Unlike the common parametric linear models (Herting et al., 2018), GAMM does not require a-priori knowledge of the relationship between the response and predictors, which enables more flexible and efficient estimation of changing patterns (Mills and Tamnes, 2014; Wood, 2017). In addition, GAMMs are well suited for the repeated measurements (e.g., our accelerated longitudinal samples from developing brains), as they account for both within-subject dependency and developmental differences among participants at the time of study enrollment (Alexander-Bloch et al., 2014; Harezlak et al., 2005). Such a GAMM was implemented using the following formula in *R* language with the **mgcv** package:

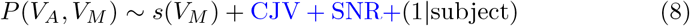

where the *s*() is a smoothing function with a fixed degree of freedom and cubic B-splines, whose number of knots is set at 5 (determined to be optimal for our data). This was set to be sufficiently large to have adequate degrees of freedom across both spline terms from fits of the model to the amygdala volume, but sufficiently small to maintain reasonable computational efficiency. The CJV and SNR were included as fixed-effect terms in the regression.

### 2.5. Modeling growth curves of human amygdala development

To fully model method-related differences in the growth curves of human amygdala volumes, we employed the following GAMM to examine age-related changes of the human amygdala by including the tracing method and its interaction with age as variables of interests:

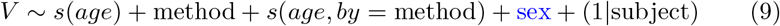

where *V* represents the amygdala volume and *s*() is a smoothing function, with a fixed degree of freedom and cubic B-splines (the number of knots = 5). Tracing method was entered as an ordinal factor (manual = 0, automatic = 1). The method term reflects the method differences in the intercept (i.e., the main effect of method). The sex term reflects the sex differences in the intercept (i.e., the main effect of sex). The first smoothing term models the slope of age for manual tracing, and the second smoothing term models the difference in the age-related slope between methods (i.e., *age*×method interaction). The *p* value associated with this term is the basis of statistical inference regarding methodological differences in developmental trajectories of bilateral amygdala volume.

To more specifically understand differences in age trajectories between methods, a set of GAMMs (see the equation 10) were proposed to detect age-related changes revealed by each method separately:

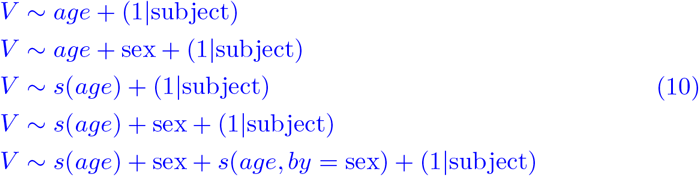

The first GAMM models the traced volume as a fixed linear age effect. As previous studies have consistently shown larger brain regions in males than in females (Herting et al., 2018), we established the second GAMM model with sex as a fixed term to assess the sex difference in the trajectory intercept. The third GAMM models the traced volume as a smoothing function of age. The fourth model is established by including sex as a fixed-effect term in the third model, as well as the fifth GAMM model including *age*×sex interaction to test the sex differences in the trajectory slope. The Akaike Information Criterion (AIC) was used to determine which model had the best fit. All fit models were tested against a null age effect model. The model chosen as the best fit model had to have the lowest AIC value and be significantly different from null. The data analyses and visualization were performed using the mgcv (Wood, 2017) and ggplot2 (Wickham, 2016) packages in R (R Core Team, 2014), respectively.

We also tested growth curves of the human amygdala by accounting for global brain features in the GAMMs. The volumes of subcortical structures are known to be related to brain size (Brown et al., 2014; Brain Development Co-operative Group, 2012; Uematsu et al., 2012). Accordingly, we included ICV as a covariate for regression control to enable the removal of individual variability that can be explained by brain size (Narvacan et al., 2017; Sawiak et al., 2018; Herting et al., 2014). Researchers have also demonstrated that the size of the amygdala often scales with the GMV (Van Petten, 2004; Rice et al., 2014). We thus accounted for brain size by controlling for the GMV in the GAMMs. We performed the analysis with CS-FS, LG-FS, *volBrain* and *FSL* derived GMV measurements and CS-FS, *volBrain* and *FSL* derived ICV measurements. To better understand the differential effects when controlling for ICV and GMV, we modeled and plotted the measurement bias as a function of either GMV or ICV. The measurement bias were quantified by the spatial overlap and false positive rate of automated segmentation with manual tracing. In addition, the growth curves were delineated for both GMV and ICV to help understanding their differential effects when controlling them for modeling amygdala’s growth.

## 3. Results

### 3.1. Measurement reliability of manually traced human amygdala

We reported almost perfect reliability of the human amygdala volumes measured by the manual tracing protocol. Specifically, as in **Table 2**, both intrarater and inter-rater reliability of the volumes for the manually traced amygdala were achieved. Inter-rater ICCs were around 0.88 with 95%CI= [0.80, 0.96] for the left amygdala, and 0.89 with 95%CI= [0.83, 0.95] for the right amygdala. Intra-rater ICCs were also almost perfect: 0.91 with 95%CI=[0.82, 0.96] for rater ZQZ while 0.95 with 95%CI= [0.89, 0.97] for rater QZ. These results confirmed that the raters’ manual tracings could be used as the gold standard or the reference for comparisons with automatic segmentation.

### 3.2. Measurement accuracy of automatically traced human amygdala

For the first-wave samples, one-way analysis of variance with repeated measures indicated significant differences in volumes of human amygdala across the five segmentation methods (left amygdala: *F* = 413.90, *p* < 0.001; right amygdala: *F* = 347.70, *p* < 0.001). Our post-hoc paired comparisons between the automatic and manual methods revealed that volumes obtained with CS-FS and LG-FS were both significantly larger than those obtained by manual tracing (CS-FS: left amygdala: *t* = 15.45, *p* < 0.001, right amygdala: *t* = 14.51, *p* < 0.001; LG-FS: left amygdala: *t* = 19.94, *p* < 0.001, right amygdala: *t* = 21.07, *p* < 0.001, respectively), which in turn were larger than those obtained with *volBrain* (left amygdala: *t* = 53.32, *p* < 0.001; right amygdala: *t* = 50.09, *p* < 0.001). The amygdala volumes obtained by manual tracing were not significantly different with those obtained by *FSL* (left amygdala: *t* = 1.75, *p* = 0.081; right amygdala: *t* = 1.36, *p* < 0.175). These findings (**Figure 1**) are reproducible for the second and third waves of samples with the exceptions of the left amygdala volumes obtained with *FSL* were significantly larger than those obtained with manual tracing (wave-2: *t* = 2.27, *p* = 0.025; wave-3: *t* = 2.10, *p* = 0.038; see Supplementary Figure S1 and S2).

As depicted in **Figure 2**, paired two-sample t-tests revealed that the spatial overlap for the left amygdala were significantly higher for *FSL* than CS-FS (*t* = 2.10, *p* = 0.037), while the spatial overlap for the right amygdala were significantly higher for CS-FS than *FSL* (*t* = 3.14, *p* = 0.002). The comparison between the spatial overlap of CS-FS and *FSL* have been inconsistently reported for the three wave data. Both CS-FS and *FSL* methods showed higher spatial overlap with manual tracing than LG-FS (left amygdala: *t* = 19.23, *p* < 0.001; right amygdala: *t* = 17.69, *p* < 0.001; left amygdala: *t* = 9.91, *p* < 0.001; right amygdala: *t* = 1.98, *p* = 0.05). The LG-FS had higher percentages of spatial overlap than *volBrain* with the manual tracing for the first-wave data (left amygdala: *t* = 14.66, *p* < 0.001; right amygdala: *t* = 19.47, *p* < 0.001). The false positive rate of amygdala were significantly different between all methods, with the *volBrain* showing the lowest value, followed by *FSL*, CS-FS and LG-FS. These findings are reproducible for the second-wave and the third-wave samples (see Supplementary Figures S1, S2). The false-negative rates were significantly higher for *volBrain* than for *FSL*, CS-FS and LG-FS segmentation of amygdala (all *ps* < 0.05). The *FSL*, CS-FS and LG-FS showed comparable false negative rates, which resulting in inconsistency comparison results for three waves of samples (**Figure 2**, Figures S1, S2). Both the left and right amygdala volumes obtained with the CS-FS, LG-FS and *volBrain* methods only showed moderate Pearson’s correlations with those obtained by the manual tracing although statistically significant (*Rs* = 0.57 − 0.62, *ps* < 0.001; **Table 3**), but did not exceed 0.8. Correlations between manual and *FSL* were obviously weaker and did not reach statistical significance (left amygdala: *r* = 0.09, *p* = 0.207; right amygdala: *r* = 0.08, *p* = 0.310). These indicated that the individual differences in volume measured by the automatic methods were not fully consistent with those measured by the manual tracing method.

**Figure 2:**
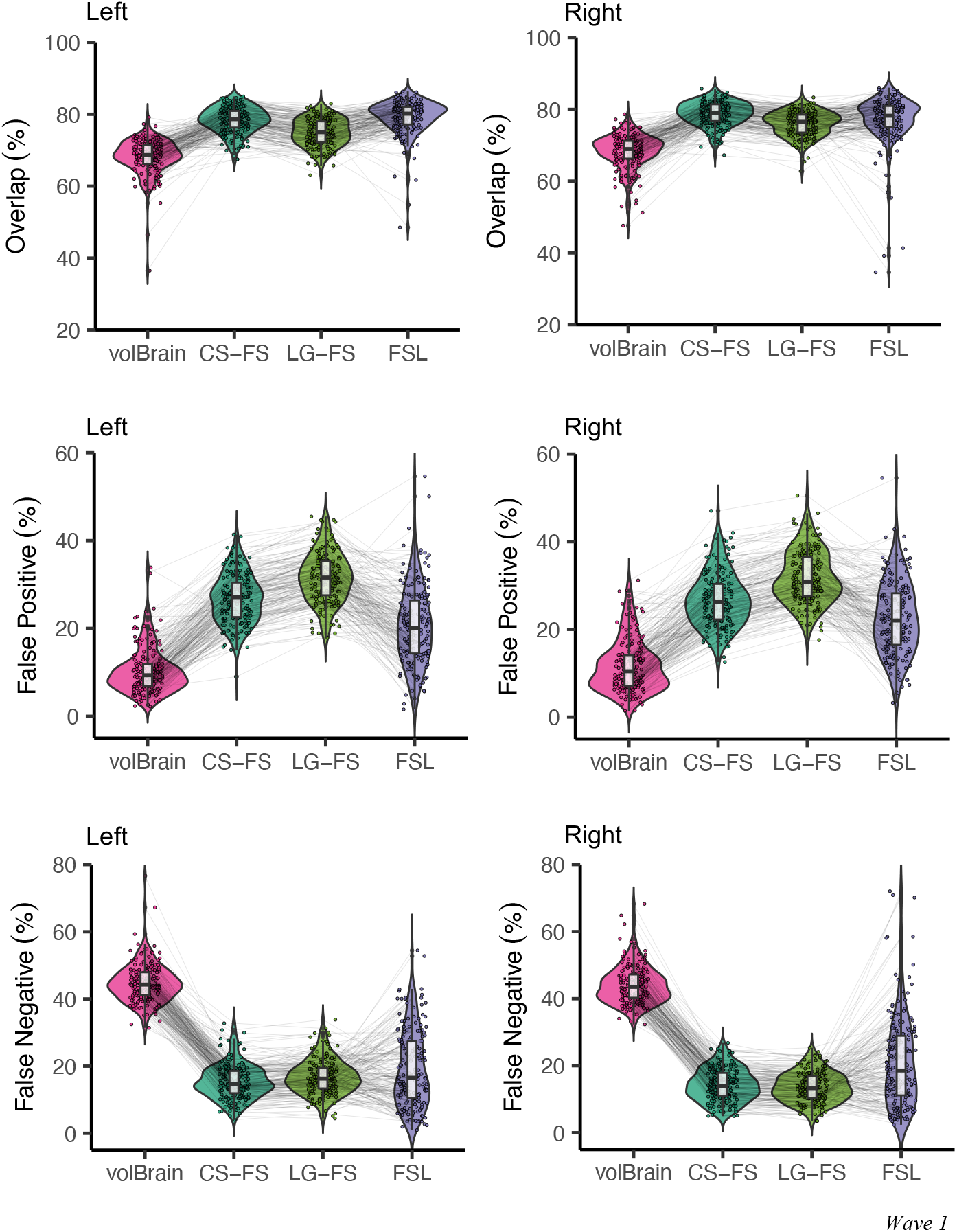
Spatial overlap, false positive rate and false negative rate for segmentation using *volBrain*, CS-FS, LG-FS and *FSL* compared to the manual “gold standard” in the CCNP wave-1 samples. Horizontal bars indicate non-significant test for difference in percent volume overlap, false-positive rate and false-negative rate. The remaining comparisons showed significant differences.

The GAMM-based regression showed that the accuracy of *volBrain* segmentation (i.e., the percentage of spatial overlap with manual tracing) increased with the amygdala size before reaching a stable accuracy with a larger volume of the amygdala (**Figure 3**; see details on the parameters for models in Supplementary **Table S1**). For CS-FS and LG-FS, the segmentation accuracy displayed a linearly significant increase pattern with the left amygdala size while a two-stage (first increase and then remain stable) pattern with the right amygdala size. The *FSL* increased and then followed by plateau with the left amygdala size while consistently remained stable with right amygdala size. In most cases of the automatic segmentation methods, a smaller amygdala structure is associated with worse segmentation accuracy after controlling image quality. These results indicated that neuroanatomical features can possibly bias the accuracy of automatic segmentation in a systematic way. In addition, the effects of image quality and motion are detectable for both *volBrain* and *FreeSurfer* segmentation accuracy while not *FSL*. The effect of more movements (higher CJV) on the segmentation accuracy of amygdala with CS-FS is significant in children during imaging while the effect of worse quality (lower SNR) on the segmentation accuracy with both *volBrain* and *FreeSurfer* were also significant (see in Supplementary Table S1).

**Figure 3:**
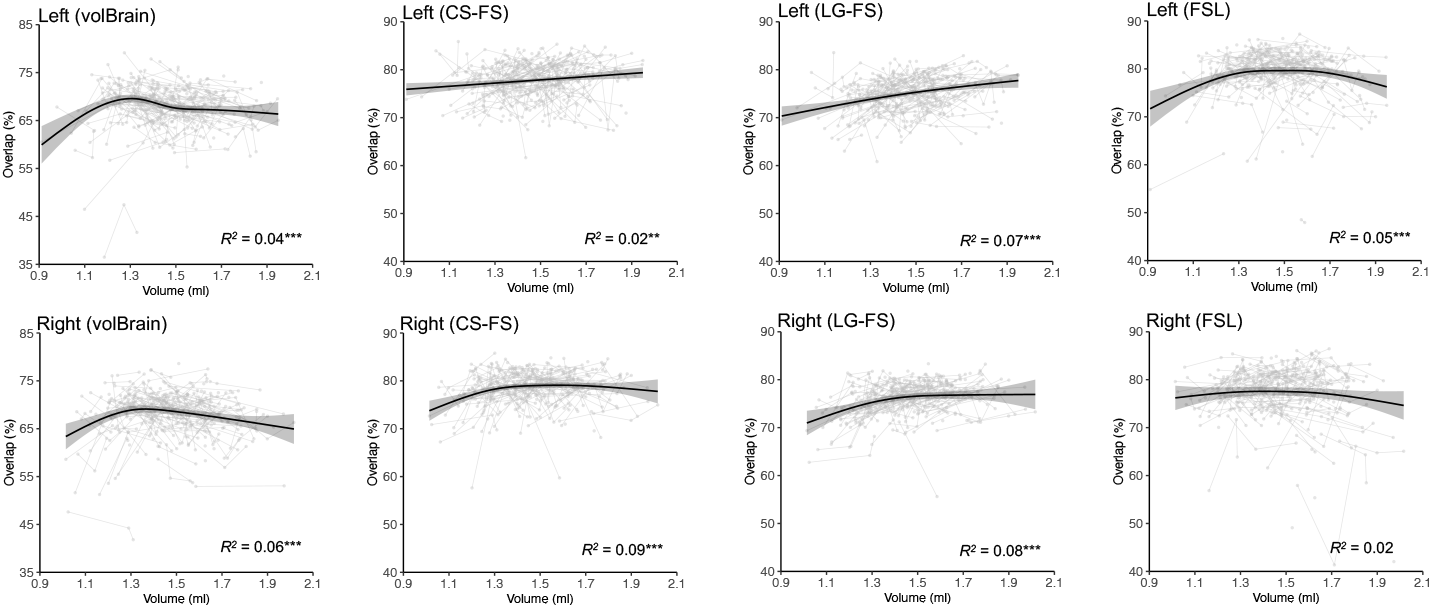
Percentage of spatial overlap of automatic methods as function of the amygdala volume. ∗*p* < 0.05; ∗ ∗ *p* < 0.01; ∗ ∗ ∗*p* < 0.001

### 3.3. Growth curves of human amygdala volume

The unified GAMM method, which includes age and interactions terms indicated that the age effects on the human amygdala were not consistent across the automatic tracing methods (**Table 4**). Specifically, these models reproduced the results of measurement accuracy for *volBrain, FreeSurfer* and *FSL* reported in the previous section. The *volBrain* produced amygdala’s age-related changes highly similar to that of manual tracing, i.e., no *age*×method interactions (all *ps >* 0.05). In contrast, the age-related amygdala changes showed discrepancies between *FreeSurfer* (including both CS-FS and LG-FS) and the manual tracing more overt than *volBrain* versus manual tracing method-differences, although no statistically significant *age*×method interaction was detected (all *ps >* 0.05). This led to a much lower explained variance using the GAMMs with CS-FS and LG-FS compared to that by the GAMMs with *volBrain* (CS-FS: left amygdala: 28% versus 80%, right amygdala: 28% versus 77%; LG-FS: left amygdala: 39% versus 80%, right amygdala: 33% versus 77%, respectively). In contrast, the statistically-significant *age*×method (*FSL* versus manual tracing) interaction was detectable (left amygdala: *p* < 0.001; right amygdala: *p* = 0.001). This indicated a significant difference in the growth rate of the bilateral amygdala volume between the *FSL* and manual segmentation.

The post-hoc method-wise GAMMs further revealed the growth patterns of the human amygdala as well as their sex differences. All best-fitting growth curves for each method determined by AIC were the smoothing-age models, which were significantly different from the null model on age effect. For all methods except *FSL*, the best models were determined by AIC as the fourth model, which included sex as a fixed effect and age adjusted for a smoothing spline function (**Table 5**), indicating no need for an interaction between age and sex. This model revealed bigger amygdalae in boys than in girls, but their growth rates did not differ by sex. Specifically, as shown in **Figure 4**, the growth curve patterns were parallel in girls and boys for both manual and automatic tracing methods although boys demonstrated larger volumes of their amygdalae than girls across the entire school age range (6-18 years old). As the reference standard, the manual tracing method revealed that the human amygdala (both left and right) exhibited linear growth during the school-age years in both boys and girls (left amygdala: *p* = 0.003; right amygdala: *p* = 0.001, respectively). The *volBrain* traced left amygdala yielded less linear growth (*p* < 0.001), and traced right amygdala yielded growth curves very similar to that established by the manual tracing method (*p* = 0.001). CS-FS tracing method produced less linear and flatter curves and not statistically significant, except for a marginal significant growth curve in the right amygdala (*p* = 0.066). This growth curve had an inverted U shape: increasing during childhood and early adolescence, and then decreasing in late adolescence (the peak age around 14.18 years old). LG-FS produced similar shape with those of CS-FS, all exhibiting somehow nonlinearity although its degree of nonlinearity is left-right flipped between the two FS segmentation methods. In addition, LG-FS detected statistical significant increases with age (left amygdala: *p* = 0.034; right amygdala: *p* = 0.022, respectively). For *FSL* method, the best models were the third model, which only included the smooth age effect. This model revealed the same pattern of trajectories for girls with boys while the bilateral amygdala exhibited two-phase growth patterns significantly different from that established by the manual tracing: robust volume increase during childhood and gradual decrease in late adolescence in left amygdala (*p* < 0.001; the peak age around 15.20 years); robust volume increase during childhood and a phase of plateau over adolescence in right amygdala (*p* < 0.001; the peak age around 16.03 years).

**Figure 4:**
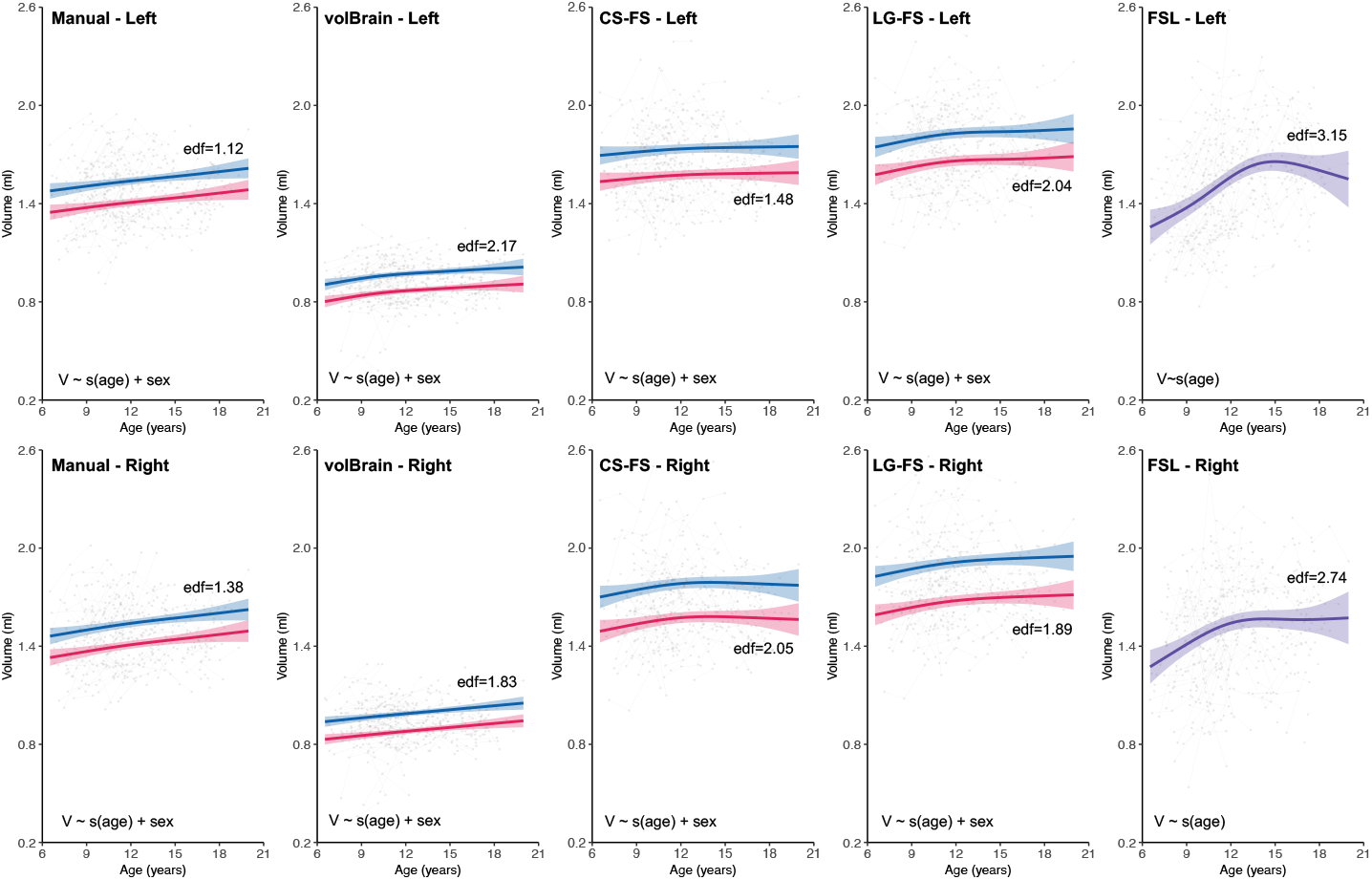
Volumetric growth curves for human amygdala traced by manual tracing, *volBrain*, CS-FS, LG-FS and *FSL*. The blue color indicates trajectories for boys while the red color indicates trajectories for girls while the purple color for each sex plotted together. The trajectories are surrounded by shaded 95% confidence intervals. Note that boys and girls showed very similar developmental trajectories with no significant age-by-sex interactions, although boys had significantly larger amygdala volumes across the school ages (all *ps* < 0.001), except *FSL* showing the same trajectories for boys and girls.

Correction for GMV abolished the significant sex differences of trajectory patterns derived by *volBrain* across the entire age range (see details on the parameters for best-fitting models in Supplementary **Table S2**). The growth patterns derived by the manual tracing method after controlling for either *vol-Brain*-estimated GMV or *FreeSurfer*-estimated GMV or *FSL*-estimated GMV remain consistent with those without the GMV corrections (**Figure 5**). After the GMV-based correction, the growth patterns derived by *volBrain*, CS-FS and LG-FS showed almost identical shapes to those obtained using the manual tracing method (**Figure 6**). This correction highly increased the reproducibility of the human amygdala growth curves across *volBrain*,CS-FS and LG-FS. The shape discrepancies between manual tracing and *FSL* derived growth curves after the GMV-based correction did not resolved. As shown in **Figure 5**, the growth patterns derived by the manual tracing method remained consistent with those without ICV corrections, but with much less statistical power: controlling for *volBrain*-estimated ICV or *FSL* estimated ICV led to much less significant age-related changes while controlling for CS-FS-estimated ICV led to no significant age-related changes (see Supplementary Table S3). Correction for ICV reduced the reproducibility of the human amygdala growth curves across the three automatic methods (**Figure 6**). The significant positive linear association with age remained for *volBrain* traced amygdala with less statistical power, even after controlling for the ICV (**Figure 6**). However, correction for ICV changed the CS-FS-derived growth curves of the amygdala volume from nonlinear (not significant) to linear decrease (significant) patterns (**Figure 6**). The growth patterns derived by the *FSL* were still inconsistent with manual tracing without ICV corrections, showing nonlinear increases (**Figure 6**).

**Figure 5:**
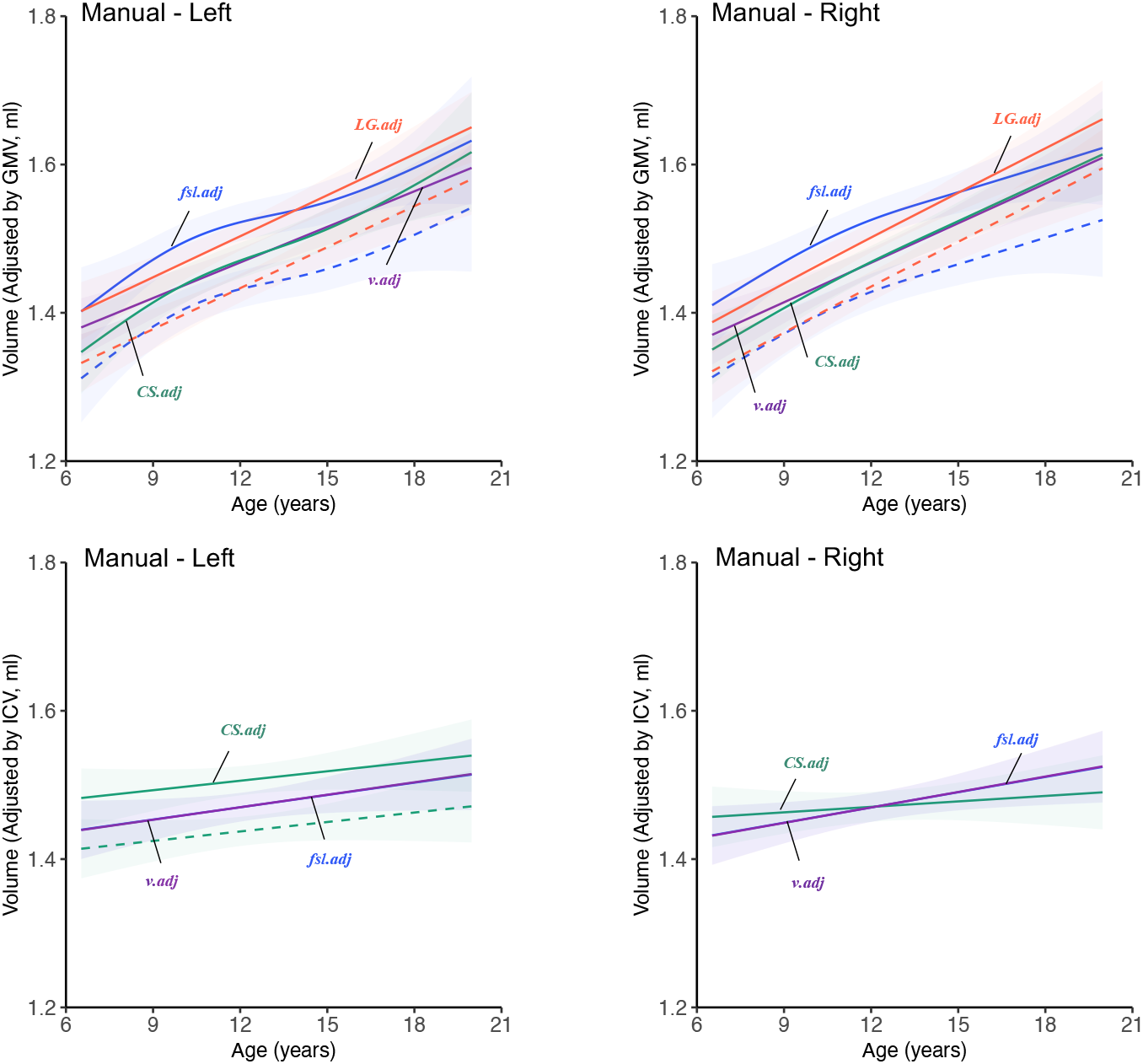
Growth curves of manually traced volume for human amygdala adjusted by GMV and ICV. Amygdala volumes are adjusted by GMV in different ways: CS.adj, adjusted by CS-FS produced GMV; LG.adj, adjusted by LG-FS produced GMV; v.adj, adjusted by *volBrain* produced GMV; fsl.adj, adjusted by FSL produced GMV. Amygdala volumes are adjusted by ICV in different ways (B): CS.adj, adjusted by CS-FS produced ICV; v.adj, adjusted by *volBrain* produced ICV; fsl.adj, adjusted by *FSL* produced ICV. solid line = male, dotted line = female. The trajectories are surrounded by shaded 95% confidence intervals.

**Figure 6:**
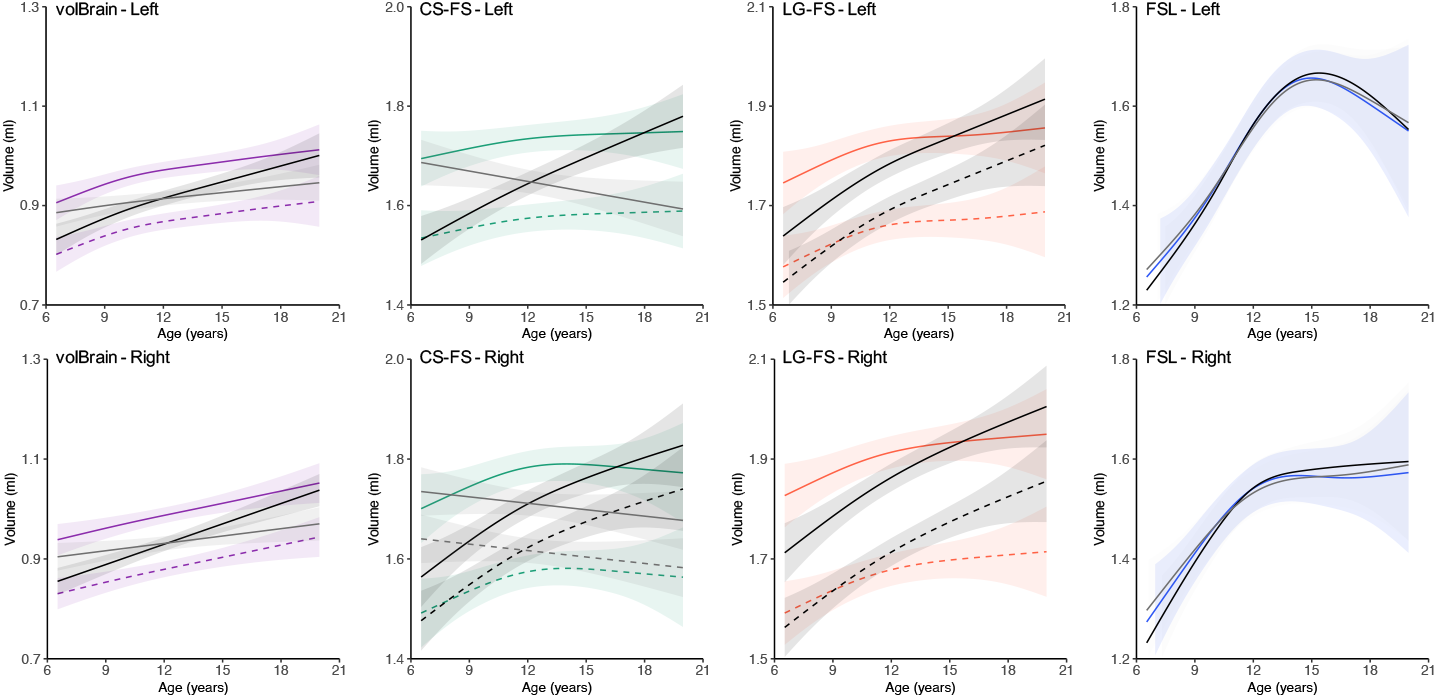
Growth curves of automatically segmented volume for human amygdala adjusted by GMV (black line) and ICV (gray line). solid line = male, dotted line = female. The trajectories are surrounded by shaded 95% confidence intervals.

## 4. Discussion

This study evaluated the performance of segmentation of the amygdala using the automatic software *volBrain*, the cross-sectional and longitudinal pipeline of *Freesurfer* and *FSL* compared to manual tracing in a longitudinal developmental sample. Importantly, we also explored how the segmentation differences could impact the growth curve modeling of the amygdala development. The findings indicated systematic differences in tracing performance across the three methods. CS-FS overestimated the volumes with more spatial overlapping with the manual tracing method, but had higher false-positive rates. LG-FS also segmented larger amygdalae than the manual method and showed smaller spatial overlaps and higher false-positive with the manual tracing than CS-FS segmentation. In contrast, *volBrain* tended to underestimate the volumes with less spatial overlap with the manual tracing method, but had lower false-positive rates. *FSL* estimated the volumes with more spatial overlapping but had inconsistency with the manual tracing. We noted that the tracing accuracy of automatic methods was worse for smaller amygdalae. Furthermore, the growth curves of the amygdala volume estimated by different methods were inconsistent. These discrepancies indicated the importance to evaluate the segmentation performance across methods, especially in a developmental sample. This study presented manual tracing of the amygdalae in a large-scale longitudinal sample and presented a systematic investigation of the method-wise variability of the growth curves of the human amygdala across school age. This variability of growth patterns of amygdala volume derived from *volBrain* and *FreeSurfer* could be normalized by adjusting for the total gray matter volume, but not adjusting for intracranial volume. The manual tracing method revealed linear growth of the amygdala in both boys and girls throughout the school-aged years, which is valuable to provide a growth norm for pediatric studies in the future.

The measurement accuracy of the amygdala volume varied across the automatic methods. *FreeSurfer* overestimated amygdala volumes (13–28%), and this overestimation has been observed in previous studies of the amygdala volume measurement by *Freesurfer* (Morey et al., 2009; *Schoemaker et al., 2016*). It is likely due to the greater variability in the definition of the amygdala boundary and liberal inclusion of voxels near this boundary (Morey et al., 2009; Schoemaker et al., 2016). The degree of overestimation observed here was greater than that reported for adults (7 − 9%) (Morey et al., 2009), but less than that reported for children aged 6–11 years (93–100%) (Schoemaker et al., 2016). In addition, FSL had comparable volume estimates with manual tracing or had slightly higher estimates than the manual method. Although the *FSL* did not excessively overestimate and underestimate amygdala volume, it showed high absolute difference percentage (17-21%) with manual tracing. The degree of volume difference observed here was also greater than that reported for adults (3− 6%) (Morey et al., 2009), but less than that reported for children aged 6–11 years (40–50%) (Schoemaker et al., 2016). Schoemaker et al.(2016) suggested it might be caused by using a standard brain template derived from adults. This may introduce greater bias when applied to a pediatric sample, in which amygdala sizes and shapes differ from adults. Another possible reason for this volume difference might be artifacts caused by more movements in children during imaging, causing a less precise differentiation and classification of amygdala structures by *FreeSurfer*. In contrast, *volBrain* underestimated the amygdala volume (35–37%) compared to the manual tracing. The underestimation may reflect the stringent inclusion of the amygdala during the segmentation by *volBrain*. This underestimation has been also observed previously, but is greater in children than for adults (3.38%) (Manjón and Coupé, 2016). *volBrain* segmentation uses manually labeled brain templates from 50 individuals with ages from 2 years old and 24-80 years old (Manjón and Coupé, 2016), which have no overlap with the age range of the current study (6-19 years old). The opposite directions of the estimation differences between *FreeSurfer* and *volBrain* methods imply, other than using unmatched templates, the vast variation in automatic extraction that may exist. Further studies are clearly warranted to explore whether the use of age-matched templates could improve the accuracy of automatic amygdala segmentation (Dong et al., 2020). Given the systematic differences in the amygdala volume between automatic and manual segmentation, it calls for caution on interpreting the results of the absolute amygdala volumes obtained by using the automatic methods in children and adolescents.

*FreeSurfer* and *FSL* exhibited more spatial volume overlap than *volBrain* with the manual tracing method. The spatial overlap (76 − 79%) observed between automatic methods (CS-FS and *FSL*) and the manual segmentation is consistent with the results reported by Morey et al. (2009). A higher overlap of *volBrain* was reported in a previous study (Manjón and Coupé, 2016), which is inconsistent with the observation in the present work. This could be related to the excessive underestimation of volume caused by the age-mismatched brain templates used by *volBrain* when segmenting amygdala for children and adolescents. In terms of spatial overlap, *FreeSurfer* and *FSL* outperformed *volBrain* for human pediatric amygdala segmentation. However, in terms of false-positive rate, *FreeSurfer* performed less than *FSL*, which in turn less than *volBrain*. The high false-positive rate of *FreeSurfer* could be an indication of its overestimation of the volume. A previous study suggested that it was due to excessive segmentation of brain structures in *FreeSurfer* by including structures and areas not part of the target structure (Næss-Schmidt et al., 2016). Correspondingly, the overestimation of *FreeSurfer* help to reduce errors of the exclusion of larger proportions of amygdalar structure, which resulting lower false-negative rates than *volBrain*. In terms of false-negative rate, *FreeSurfer* performed best among these automatic methods. Although few studies have explored the performance of *volBrain* on human amygdala segmentation in terms of the false-positive rates and false-negative rates, similar performance results have been shown for the automatic segmentation of the hippocampus and thalamic volume (Næss-Schmidt et al., 2016). According to inter-individual differences in segmented amygdala volumes, the two automatic methods only demonstrated moderate correlation with the manual segmentation, while *FSL* was not significantly correlated with manual tracing. These are consistent with previous work (Morey et al., 2009; Grimm et al., 2015), implying its potential challenge for reliable measurements of their growth curves. Though, LG-FS showed slightly lower (not statistically significant) false-negative rates than the CS-FS segmentation, which outperformed LG-FS in terms of volume difference, spatial overlap and false positive rate metrics. The possible bias by matching head sizes across all the time points in children caused the worse accuracy of LG-FS than CS-FS. Overall, the CS-FS, *volBrain* and *FSL* methods have advantages and disadvantages for the assessment of amygdala volume. The complex amygdala structure adds difficulty to reliably and validly estimate its volume. It’s a trade-off to choose which method should be used, requiring careful evaluation, and also demonstrates which facet of the automatic methods should be further improved in the future.

In this study, we found that the automatic segmentation performed worse in smaller amygdalae in developmental neuroimaging studies of children and adolescents. After controlling the quality of scan, the segmentation accuracy increased with amygdala volume, and then remained stable when the amygdala has reached a large enough size (around 1.3-1.4 ml). Previous studies have found that smaller brain structures were associated with greater automatic segmentation errors due to their sizes and shapes differing from adults (Schoemaker et al., 2016; Biffen et al., 2020; Sánchez-Benavides et al., 2010). Our results are consistent with that neuro-anatomical and geometric features could systematically influence the accuracy of their automatic segmentation. This bias is likely less problematic in adults, whose structures are commonly larger than in children. Poor scan quality caused less precise differentiation and classification of amygdala structures when using *FreeSurfer* and *volBrain* (see Supplementary TableS1). However, after controlling the quality of scans, the segmentation accuracy was still improved when the amygdala volume increased, and then remained stable when the amygdala reached a large enough size. The patterns of the spatial overlap function were highly similar between the two models (i.e., with/without controlling CJV and SNR). Therefore, we considered the nonlinearity presented in **Figure 3** is more likely attributed to the use of age-unmatched brain templates and the anatomical complexity of human amygdala. Previous studies on the amygdala have reported that the human amygdala could undergo the significant and complex deformation during childhood and adolescence (Schoemaker et al., 2016). The adult templates used in the automatic protocols mismatched with the current developing samples, especially with the young children, more likely causing poor segmentation results of young amygdalae. The human amygdala has been widely investigated in pediatric studies and associated with many developmental disorders such as autism (Mosconi et al., 2009; Schumann et al., 2009, 2004) and anxiety disorder (De Bellis et al., 2000; Hill et al., 2010; Milham et al., 2005). Our findings further highlighted the importance of using age-matched template and improving the measurement accuracy of automatic segmentation for developing individuals. We argue that, in the current stage, manual tracing should be given priority for amygdala volume estimation in pediatric research. In the future, eliminating the bias in automatic segmentation methods will be of great importance.

Although the statistical models indicated that the systematic differences in amygdala volume exhibited moderately marginal effects on growth curve modeling between the *FSL* and manual segmentation, our post-hoc growth chart analyses demonstrated remarkable discrepancies in age-related changes of the human amygdala across development. As the ‘gold standard’, manually traced amygdala volumes exhibited linear growth patterns without sex differences in growth rate. This is completely consistent with the patterns validated by the manual tracing method for amygdala growth from youth to adulthood in the macaque monkey (Schumann et al., 2019). Most previous studies of amygdala development in children and adolescents have been based on automatic segmentations (Wierenga et al., 2014; Goddings et al., 2014; Herting et al., 2018; Uematsu et al., 2012) while manual segmentation has been used in only two studies (Giedd et al., 1996; Merke et al., 2003). We noted that the developmental patterns of the amygdala have been inconsistent across these studies between automatic and manual methods. The growth patterns we detected by manual tracing were generally consistent with that by Giedd et al. (1996) and Merke et al. (2003) although they observed volume increases only in boys, but not in girls. In our study, the amygdala volumes grew in both boys and girls along highly similar trajectories. Such distinction may be an indication of the difference in scanning field strengths (3T versus 1.5T). Higher-resolution MRI enabled us to detect subtle changes in the human amygdala volume. Regarding automatic segmentation, previous studies generated amygdala growth curves with inverted U shapes from childhood to adolescence with peaks around 12–15 years old (Wierenga et al., 2014; Goddings et al., 2014; Herting et al., 2018; Uematsu et al., 2012). These were similar to our findings based on CS-FS and *FSL* segmentation, which showed a nonlinear trend of growth, especially for the right amygdala, with an inverted U-shaped trajectory (the volume peak at 14.18 years old). LG-FS produced similar shape with those of CS-FS, all exhibiting somehow nonlinearity although its degree of nonlinearity is left-right flipped between the two FS segmentation methods (**Figure 4**). Although the LG-FS detected the statistical significance of amygdala’s age-related increases, it achieved by overestimating amygdala volume more than CS-FS and sacrificing its segmentation accuracy. Therefore, only in the case of exploring the growth trend of the amygdala, LG-FS is preferable to CS-FS. *volBrain* segmentation yielded growth curves most similar to that obtained by the manual tracing for the amygdala development. *volBrain* seems to have less error modeling growth curves than *FreeSurfer*. However, given limited studies using *volBrain* to investigate amygdala development in children and adolescents, it is hard to compare our results with others directly.

The growth curves between the automatic methods (except for *FSL*) and manual tracing became similar when we adjusted amygdala volumes by the total gray matter volume rather than intracranial volume. This may reflect the reduction in the bias related to the amygdala size in automatic segmentation as mentioned above correcting the amygdala volume. As shown in Figure S7, the GMV derived with all methods (except for *FSL*) decreased as growing. The inclusion of GMV as a covariate in regression of amygdala volume on age could explain some variances of the amygdalar volume decreases (ei.ge., the decreasing part of the inverted-U shapes), resulting in the changes of developmental patterns from weak linearity to strong linearity as we demonstrated. We also found that a smaller GMV was associated with worse performance of automatic amygdala segmentation (except for *FSL*) but remain stable for large enough GMVs (Figure S4). As the GMV decreased with age, the measurement bias for younger children could be removed more than older children by controlling for GMV. This further strengthened the linearity of the developmental patterns of *volBrain* and *FreeSurfer*. In contrast, the measurement bias of *FSL* was negatively related to GMV, leading to more corrections of the measurement bias for older participants and thus the aggravated decreases. As shown in Figure S8, the ICV obtained with the three methods all increased when growing. The inclusion of ICV as a covariate in regression could explain some variances of age-related increases of amygdala volume. *volBrain* demonstrated a segmentation bias associated with ICV (Figure S5), indicating a smaller brain associated with worse amygdala segmentation but remain stable for big enough brains. This might cause the changes of developmental patterns of *volBrain*-derived amygdala from strong linearity to weak linearity. However, CS-FS did not demonstrate such an ICV-related segmentation bias (Figure S5) but higher false positive rates of segmentation associated with larger brains (Figure S6). After controlling ICV, the amygdala volumes of the smaller brains were corrected little, while the volume estimates of the larger brains were corrected more. This might explain that the changing pattern of CS-FS derived growth curves from increasing with age to decreasing with age. These results suggest that controlling for the gray matter volume improved the accuracy of curve-fitting on the *volBrain* and *FreeSurfer* of amygdala from childhood to adolescence.

Accurate delineation of the development of the human amygdala is fundamentally important by providing neuroimaging biomarkers for various developmental disorders (DiMartino et al., 2014; Zuo, 2020; Holla et al., 2020). Our findings present an unaddressed bias and challenge for charting the growth of the human amygdala across school-age children and adolescents-the growth curve modeling was highly dependent on the segmentation method. The methodological differences may contribute to the inconsistencies among previous findings regarding the patterns of amygdala development during childhood and adolescence (Wierenga et al., 2018; Uematsu et al., 2012; Albaugh et al., 2017; Herting et al., 2018). Given the inconsistency, we give researchers working on the amygdala of children and adolescents some suggestions. 1) Manually tracing the amygdala if possible. This seems affordable but prohibitive for very large-scale MRI datasets, considering it takes 90 minutes to manually tracing an amygdala and will take 3 months to manually trace 500 amygdalae. A practical solution would be distributing the mission to a tracing team achieved high within-operator and between-operator reliability of the tracing operation by an established training protocol. 2) Checking and correcting the automatic segmentation of the amygdala by a trained professional to improve the accuracy and save the effort if the manual segmentation is not feasible. This would likely lead to significantly improved accuracy and time cost of the tracing segmentation although need further investigations in future. In addition, a very promising solution is to train a computational segmentation tool by integrating the knowledge aggregated from big data of the manually segmented amygdalae by using some novel methods (e.g., the deep learning algorithms). 3) Using age-matched brain templates for automatic segmentation (Dong et al., 2020). 4) Using high-resolution MRI protocol on scanning the amygdala. 5) Adjusting the amygdala volume by total gray matter volume when conducting statistical analysis. 6) Comparing and interpreting previous findings cautiously using different segmentation methods than the study proposed, particularly for the smaller amygdalae (e.g., younger children and females). To facilitate the use of the growth curves we developed for human amygdala development at school age, we generated their charts and made them publicly available to the community (LINK TO BE ADDED after a final publication).

Our study has some limitations that should be noted. First, the age span of our sample might not be sufficient for examining the full range of development of the human amygdala from childhood, adolescence and into young adulthood. While the previous work in the macaque monkey revealed the linear pattern of amygdala growth from youth to adulthood (Schumann et al., 2019), further work would benefit from the extension of the age span into adulthood for direct growth assessments in human in future. Second, we did not investigate the measurement reliability across different versions of automatic segmentation tools, which has been shown remarkable influences on the brain segmentation (Gronenschild et al., 2012). This factor should be carefully evaluated by using different versions of these tools to model amygdala growth. Finally, we only examined the overall volume measurement of the human amygdala. In the future, we will employ more local and shape measurements (Li et al., 2012; Roshchupkin et al., 2016) for investigating more details of human amygdala growth. To provide more efficient and accurate tracing of the pediatric amygdala, we also plan to develop an automatic algorithm based upon the manually traced samples using more advanced methods such as deep learning (Ataloglou et al., 2019).

## 5. Conclusion

By manually tracing a large-sample pediatric MRI dataset from the accelerated longitudinal cohort, we charted the growth of human amygdala across school age. We identified measurement biases for the automatic amygdala segmentation methods and their impacts on modeling growth curves of the amygdala volumes from childhood to adolescence. There is considerable room for the methodological improvement on next generation tools of automatic segmentation to achieve more accurately tracing of the human amygdala during development. Our work provides not only a practical guideline for future studies on amygdala in children and adolescents but also its growth standard resources for translational and educational applications. This can be implemented with normative modeling (Holla et al., 2020; Marquand et al., 2016) for individualised assessments on typical or atypical development as well as their associations with behavioral performance and school achievement.

## Acknowledgments

We thank all families participating in the Chinese Color Nest Program and all the support from schools and communities. This work was supported in part by the Key-Area Research and Development Program of Guangdong Province (2019B030335001), the Start-up Funds for Leading Talents at Beijing Normal University, the National Basic Science Data Center “Chinese Data-sharing Warehouse for In-vivo Imaging Brain” (NBSDC-DB-15), the Beijing Municipal Science and Technology Commission (Z161100002616023, Z181100001518003), the CAS-NWO Programme (153111KYSB20160020), the National Natural Science Foundation of China (81220108014), the Guangxi BaGui Scholarship (201621) and National Basic Research (973) Program (2015CB351702). Finally, we would like to thank Tonya White, MD, PhD and Neda Jahanshad, PhD for their comments, suggestions, and for proofing the article for English language.

